# Hierarchical Extended Linkage Method (HELM)’s Deep Dive into Hybrid Clustering Strategies

**DOI:** 10.1101/2025.03.05.641742

**Authors:** Lexin Chen, Jherome Brylle Woody Santos, Jokent Gaza, Alberto Perez, Ramón Alain Miranda-Quintana

## Abstract

Clustering remains a key tool in the analysis of molecular dynamics (MD) simulations, from the preparation of kinetic models to the study of mechanistic pathways and structural determination. It is no surprise then that multiple algorithms are currently used in the MD community, with *k*-means and hierarchical approaches being arguably the two most popular approaches. The former is very attractive from a purely computational point of view, demanding minimal memory and time resources, but at the price of being able to partition the data in very restrictive ways. Hierarchical strategies, on the other hand, can generate arbitrary partitions, but with steep memory and time requirements due to their need to build a pairwise distance matrix for all the considered conformations/frames. Here we propose a new hybrid paradigm, the Hierarchical Extended Linkage Method (HELM), that retains the efficiency of *k*-means while incorporating the flexibility of hierarchical methods. The key ingredient is the use of *n*-ary difference functions as a way to stabilize the *k*-means results and efficiently build the hierarchy of subsets. We showcase the applicability of this strategy over protein-DNA and protein folding studies, including the complete analysis of simulations with over 1.5 million frames. HELM is freely available in our MDANCE clustering package.

## 1. INTRODUCTION

Between 20-40% of current academic supercomputer time goes to molecular dynamics (MD) simulations, which provide a glimpse of the life of biomolecules as they access different configurations along time^1^. Multiple replicates on timescales exceeding the microsecond timescale are now common to sample biomolecule’s frustrated energy landscape. Together with improvements in computer power and force field accuracy leads to ever increasing molecular ensembles^1–3^. Typically, these ensembles are analyzed based on observed biomolecular states, defined based on configurations that cluster together along some similarity measure. Each cluster’s population and representative structure, often the centroid of such clusters, are used to represent and characterize the ensemble^4^. However, improvements in how trajectories are clustered have not kept pace with the increase in ensemble size. In many instances this leads to clustering on only a small fraction of the dataset, missing potentially important states.

Clustering methods must balance two often conflicting requirements: computational complexity and flexibility to partition the data. For example, methods like *k*-means^5,6^ are very popular mostly because they are time- and memory-efficient, even if they can have some well-known drawbacks. The most common reasons mentioned in the literature to avoid using *k*-means are usually the difficulty in picking a sensible *k* value and the potential instability in the final assignments^7,8^. However, the former can be addressed by carefully following the global or local behavior of clustering quality metrics, like the Davies-Bouldin (DBI)^9^, Dunn^10,11^, Calinski-Harabasz (CHI)^12^, or silhouette indices^13^, while the latter can be tackled with deterministic seed selection methods like the N-Ary Natural Initialization (NANI)^14^ protocol. Here, we want to focus on another issue instead: *k*-means can only produce very restricted clusters. At any given iteration, points are assigned to the closest of *k* centroids, which means that the data can only be partitioned in convex shapes (a “Voronoi tessellation”, see Fig. 1)^14,15^. This is not a great problem when one is only interested in a bird’s-eye view of the data, since *k*-means will still partition most of it correctly, but some fine details will inevitably be lost.

**Figure 1:**
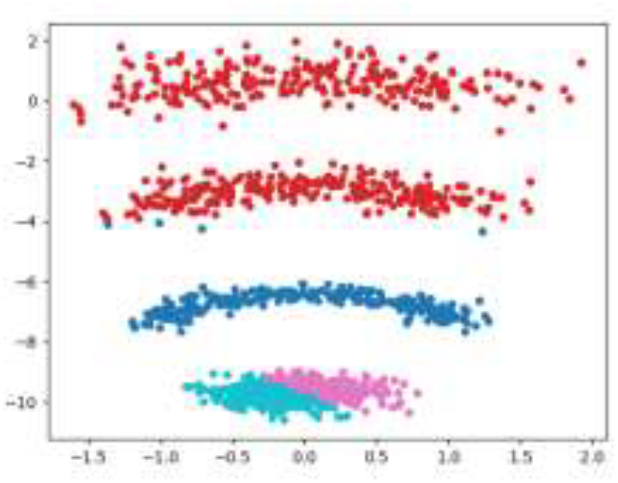
*k*-means clustering of a model 2D system with non-convex clusters.

On the other hand, hierarchical agglomerative clustering (HAC) approaches have been praised for their versatility, since they can in principle organize the data in clusters with arbitrary shapes. All HAC methods are based on the same idea: given a collection of sets, the algorithm creates a new cluster by combining the elements of the two closest sets^16^. This simple recipe makes HAC very attractive due to its built-in interpretability, since we can easily follow the process of construction of the clusters throughout the hierarchical ladder. However, since the distances between clusters depend on relations between pairs of points, HAC can quickly become computationally intractable^17^. As shown in Fig. 2, HAC methods scale quadratically in both time and memory due to the need to pre-compute all the pairwise root-mean-square deviations (RSMDs) between the conformations/frames in the simulation.

**Figure 2:**
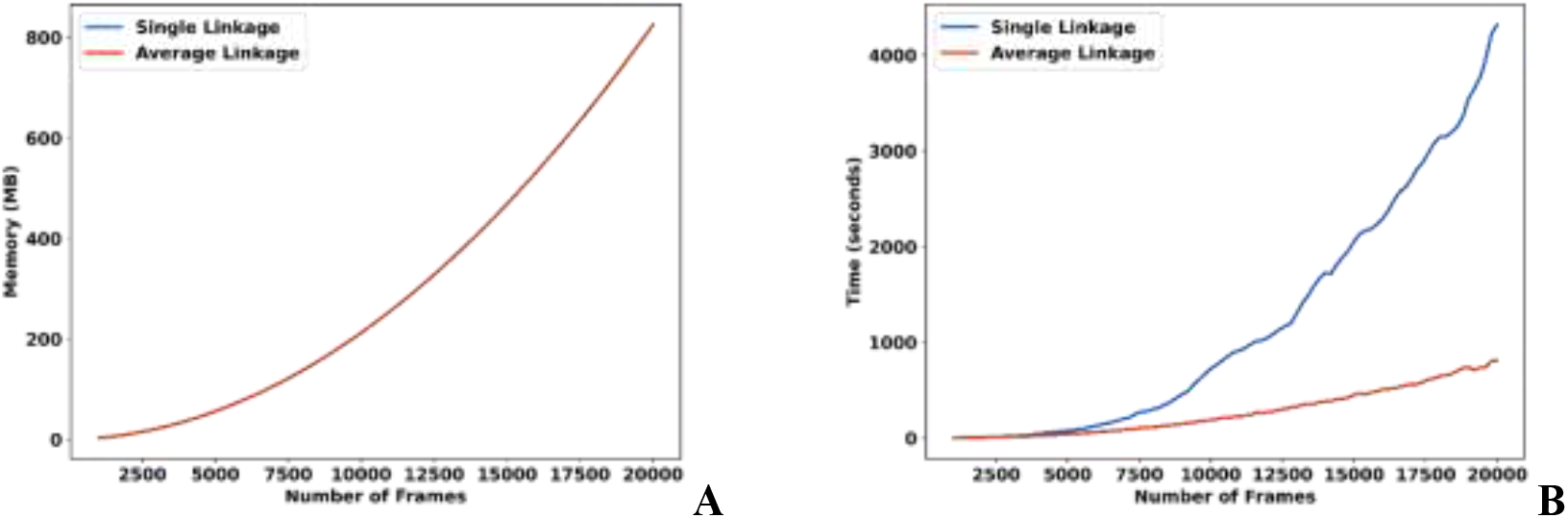
(A) Memory and (B) time dependency of single (blue) and average (red) linkage hierarchical agglomerative clustering with respect to the number of frames (frames were taken from the HP35 headpiece simulation).

From this short discussion it seems apparent that *k*-means and HAC have complementary advantages and disadvantages, so it would certainly be desirable to have new families of “hybrid” clustering methods that could potentiate the former, while ameliorating the latter. This is precisely what we propose with the Hierarchical Extended Linkage Method (HELM) approach presented in this manuscript. We leverage the recently introduced *n*-ary similarity idea to seamlessly unite *k*-means and HAC without compromising the computational cost or flexibility of these methods. The upcoming section describes the *n*-ary methodology and how it can be used to blend *k*-means and HAC. We then present three systems representative of biological challenges of interest (protein G, protein-DNA binding, and the HP35 headpiece) used to test the different HELM variants. Our results confirm that HELM greatly reduces the computational burden of traditional HAC, bringing it down to *k*-means’ levels of efficiency. We also show that HELM’s results are quite consistent with respect to the choice of the method’s conditions. HELM is publicly available as part of the MDANCE^18^ package: https://github.com/mqcomplab/MDANCE.

## 2. THEORY: *n*-ary Comparisons and HELM

Majority of MD studies uses the RMSD as the de facto way to quantify the “separation” between different conformations^19–21^. That is, if the *i*th frame, *F* ^*(i*)^, is represented using coordinates 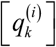, then the RMSD between frames *i* and *j* is given by:

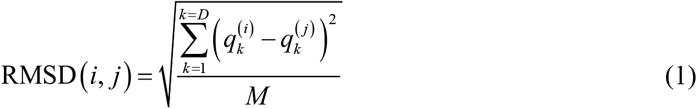

where *M* is the number of atoms and *D* is the number of coordinates used to represent the conformations.

A key problem with this expression is that calculating the average RMSD over a set of *N* points/frames requires O(*N*^2^) calculations, which is impractical for all but the smallest sets. This is not a unique feature of the RMSD, since every traditional similarity or distance metric defined over two points shares this same expensive scaling. In the field of cheminformatics, this was solved with the introduction of extended^22,23^ and instant^24^ similarity indices that can take as inputs any number of objects at the same time (they are *n*-ary functions), which allow calculating averages of similarity indices over billion-sized sets in linear time^25^. In the case of MD simulations, we have recently advocated for the use of the mean squared deviation (MSD), as a way to mimic the RMSD behavior, but with O(*N*) scaling^14,26^. Using the notation as above we can define:

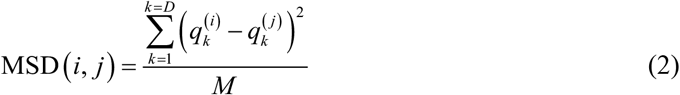

The crucial difference comes when we try to calculate the average of the MSD values over *N* frames:

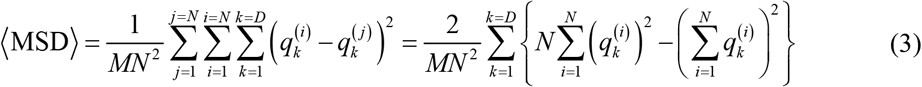

The last two summations over the frame (*i*) index make it evident that no pairwise calculations are actually required to determine the average MSD.

The first application of the average MSD is obvious: it provides a direct proxy to quantify the diversity of a set of conformations. That is, between two sets A and B, if MSD(A) > MSD(B) we can say that the conformations in B are more tightly packed. This also leads to two key results:

1. Diversity selection^23^: from a given pool of conformations, we can find a maximally diverse subset by picking those conformations that maximize the MSD.
2. Ranking conformations: The complementary MSD (cMSD) of a frame is defined as the MSD of the set after removing the frame. The relative magnitude of the cMSD in a set indicates which conformations are more representative (higher cMSD) or which are outliers (lower cMSD).

These are the key ingredients of the *k*-means NANI algorithm, which we proposed as a way to improve the selection of the initial seeds^14^. The final goal is to pick potential centroids that belong to dense regions of the data, while being separated from each other. The first task is guaranteed by selecting the top 5-10% of frames with highest cMSD values. Then, from that initial pool (and starting from the medoid of the set), the remaining *k* – 1 candidates are sampled using the diversity picking algorithm.

As noted in the Introduction, HAC methods differ on how they quantify the separation between sets (how the “linkage” is being performed). For example, two popular HAC linkage alternatives, single and average, define the distance between sets A and B as:

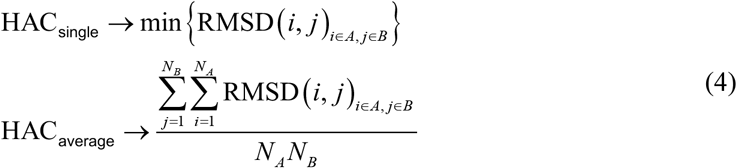

where *N*_*A*_, *N*_*B*_ are the number of frames in sets A and B, respectively. Note that to calculate both these linkages we need evaluate all the RMSDs between points assigned to different clusters. For this, we need to keep track of the individual conformations in each set. As we will see shortly, the MSD not only provides a natural way to define other linkages, but it does so without the burden of having to keep the individual information of the conformations.

It is possible to define (infinitely) many inter-cluster distance criteria based on *n*-ary functions. Here, we choose to focus on arguably the two simplest ones: *intra*- and *inter*-set dissimilarities (from now on, and for the sake of brevity, only referred to as intra and inter):

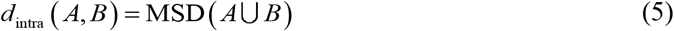

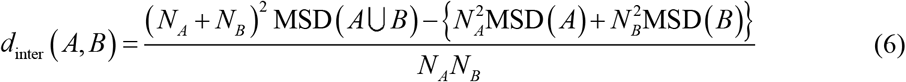

*intra* just quantifies the average MSD of the union of all the frames in A and B. *inter*, on the other hand, closely resembles the classical average HAC criterion, but instead of calculating the average RSMD separations it provides the average MSD between conformations in A and B. As remarked before, a key advantage of this approach is that we do not need to store all the individual frames in each cluster. In this regard, MSD takes inspiration from the cluster features used in the BIRCH clustering, since for a given cluster A we only need to store one number: *N*_*A*_, and two vectors: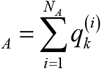 and 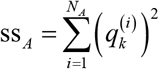, the linear sum of all the coordinates and the sum of squares of all the coordinates, respectively. The critical point is to realize that when two clusters are merged, these quantities can be trivially updated according to: *N*_*A*∪*B*_ = *N*_*A*_ + *N*_*B*_, ls_*A*∪*B*_, ls_*A*_ + ls_*B*_, ss _*A*∪*b* ⋃ *B*_ = ss _*A*_ + ss_*B*_.

With this, we now have all the pieces in place to introduce the HELM framework (see Fig. 3).

**Figure 3:**
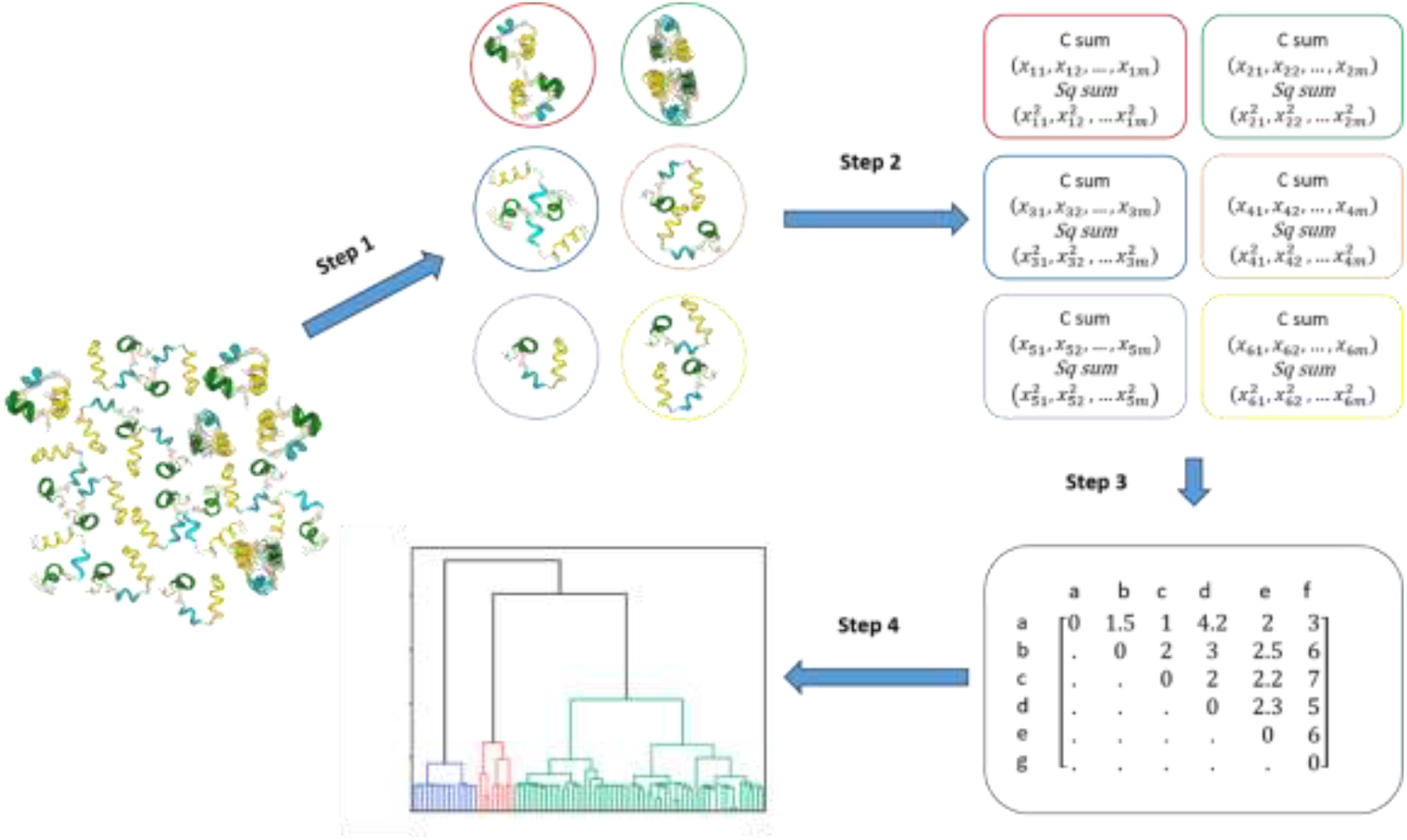
HELM workflow.

### Step 1

Perform a *k*-means NANI pre-clustering of the MD simulation: HAC studies begin with all the conformations being assigned to separated clusters (so one usually starts with ∼10^3^-10^6^ clusters), which are then combined one by one until just a few tenths are left. However, to do this one needs also in the order of 10^3^-10^6^ steps, which are mostly spent combining singletons^17^. The *k*-means step greatly accelerates this process, by providing a more convenient starting point to the hierarchical procedure. In a traditional *k*-means analysis, determining a precise *k* value is paramount, however, since NANI is just needed to pre-organize the data, we now only need to pick a *k* that we expect to be bigger than the actual number of final clusters in the set. This is a much more manageable task and only requires a very superficial level knowledge of the system. We anticipate that in most applications, the value of *k* in the 30-60 range will be more than enough. (Below we discuss the impact of choosing *k* = 30, 60, 100). Finally, the choice of NANI is motivated by the need to have reproducible results. Other *k*-means implementations can provide wildly different results from one run to another, which would severely compromise the stability of the ulterior hierarchical steps. Only NANI can guarantee a robust, fully deterministic, starting point. This step scales as O(*N*).

### Step 2

Condensing the NANI clusters: To proceed forward from the *k* NANI clusters we just need to calculate the number of frames in each of them, as well as the linear sum and sum of squares of their coordinates. This simplified representation is the key to speeding-up the hierarchical process. Now, instead of dealing with *N* ∼ 10^3^-10^6^ conformations, we only need to focus on *k* ∼ 10 starting points. This step is scaled as O(*N*)

### Step 3

Calculate the *k*x*k* matrix of inter-cluster separations: For this, we either use the intra or inter criteria (Eqs. (5) or (6)). Since we do not need information about all the individual frames in each cluster, this step scales as O(*k*^2^), so is O(*1*) with respect to the original number of frames. <u>Step 4</u>: Combine the k clusters following a hierarchical procedure: This will give the freedom to the NANI clusters to be combined in ways that are out of reach for any *k*-means method. At this point, since we start from a *k*x*k* matrix, the user can decide to build the hierarchy in a variety of ways, either: a) continuing to combine the clusters using the intra criterion; b) continuing to combine the clusters using the inter criterion; c) using the *k*x*k* matrix as the input to a traditional HAC procedure using single, average, Ward linkage^27^, etc. At worst, this is also an O(*k*^2^) step, but careful implementation of the hierarchical procedure can bring it down to O(*k*log*k*), but yet again, is O(*1*) on *N*.

Overall, the whole process scales as O(*N*), with NANI being the most computationally demanding step, since *N* >> *k*. Notice that Step 4 effectively “refines” the *k*-means clusters but also provides the information of how the clusters are related via the hierarchical tree. A nice feature of HELM is that after the *k*-means step we can analyze those clusters and decide whether they should be included in the posterior hierarchy or not. *k*-means has no notion about the noise present in the data, and every single point will be assigned to a cluster. However, if we study both the population and MSD of the *k*-means clusters we can determine if they contain mostly singletons and/or very disjoint conformations. Those clusters correspond to noisy, low-density, regions and can then be excluded from the HAC step.

## 3. SYSTEMS and COMPUTATIONAL DETAILS

### MELD Simulations

MELD (Modelling Employing Limited Data)^28,29^ is an integrative structural biology tool that combines noisy and ambiguous experimental datasets with atomistic molecular dynamics simulations through Bayesian inference, *p*(*x*|*D*) ∝ *p*(*D*|*x*) · *p*(*x*). The prior distribution, *p*(*x*), is the original Boltzmann distribution of the conformations *x* as under a given force field. Using the likelihood function *p*(*D*|*x*) ∝ exp[−*βE*_*r*_(*x*)] to represent the agreement between the sampled structure (*x*) and some subset of the data (*D*), we update the prior to obtain the posterior distribution *p*(*x*|*D*), which represents the probability of a conformation *x* given data *D*.

In MELD simulations, data are converted into flat-bottom potential restraints organized into groups (sets of restraints) and collections (sets of groups). The likelihood function allows the exploration of a subset of these restraints at each organization level given the total restraint energy, *E*_*r*_(*x*), as to mimic the potential ambiguousness of the original data^30^. The MELD potential is sampled by H,T-REMD^31^ (Hamiltonian, Temperature Replica Exchange Molecular Dynamics), which scales both the temperature and the restraint strengths along a 30 replica ladder.

In the absence of data, general directives such as “proteins form hydrophobic cores” and “β-strands can pair” can be used to guide the folding of small proteins or formation of protein-DNA complexes^32,33^. The protein G dataset was obtained from a MELD folding simulation of wild-type protein G (PDB: 3GB1) starting from an extended structure. Strand pairing was promoted by introducing distance restraints between β-strand residues separated by at least four residues. In the protein-DNA system, distance restraints between the Cα atoms of the bZIP (PDB: 1DH3) protein dimer’s binding region and the N1 atoms of the purine nucleotides in the DNA drive the binding process. This protocol assumes knowledge of the structure of the protein (i.e. the bound conformation), the DNA binding site, and the DNA-binding domain of the protein. The DNA structure follows the canonical B-form of the DNA sequence found in the PDB.

Both protein G and protein-DNA simulations were performed using the OpenMM^34^ engine with GBneck2^35,36^ implicit solvent model for protein G and GBNeck2Nu for the protein-DNA system. Proteins were defined by the AMBER ff14SBside force field^37^, and the DNA by parmBSC1^38,39^. Further details on the restraint scaling across replicas and the choice of distance restraints are available in the original MELD publications.

### Protein G

For the protein G dataset^40,41^, six distinct clusters were generated from MELD simulations, along with one noisy set, to assess the robustness of each linkage criterion. These six clusters were then concatenated into a single trajectory for further analysis, with the cluster labels already known. The single-reference alignment was performed after aligning to the first frame. For the alignment clustering, the atom selection included the alpha carbons of residues 1 to 56. A total of 1924 frames were used in this process.

### Protein-DNA Binding

Similar to Protein G dataset, six distinct clusters were generated from MELD simulation and later concatenated into a single trajectory for further analysis^32^. All frames were aligned to the heavy atoms of the DNA (residues 111 to 152) with reference to the first frame. For the clustering, the atom selection included the protein-DNA interface, specifically residues 1-27 and 57-81 of the protein, and residues 111-152 of the DNA. A total of 1517 frames were used. **HP35** The 35-amino acid chicken HP35 headpiece protein has three helical segments: helix 1 (green) consists of residues 4 to 10, helix 2 (blue) includes residues 15 to 19, and helix 3 (yellow) comprises residues 23 to 32^42^. An analysis was conducted on a 305 μs all-atom simulation of the Nle/Nle mutant of the C-terminal subdomain of the HP35 headpiece (HP35) from D. E. Shaw Research^43^. The simulation, which ran at 360 K, included 1.52 million frames with a 200 ps separation between frames. The frames were aligned to the 5000th frame, and frames before this were discarded as part of the relaxation phase. The backbone atom selection included the N atom of residue 1, Cα, C, N atoms of residues 2 through 34, and the N atom of residue 35, following the approach of previous studies on HP35. A sieve was applied to every 10th frame, resulting in approximately 152,000 frames used for clustering.

### These systems are represented in Fig. 4

A critical part of any clustering study is how to quantify the appropriateness of the final partition. This is particularly challenging since in practical applications we do not knoew beforehand which subsets optimally divide the data (if we knew this, then the actual clustering would become irrelevant!). To help with this we considered two clustering quality indicators: the Davies-Bouldin and Calinski-Harabasz indices (DBI and CHI, respectively). In short, they both measure how tightly packed the clusters are, and how well-separated they are. The traditional way to carry on this analysis just looks at the global levels of these indices, keeping in mind that optimum partitions should correspond to lower DBI values and higher CHI values. So, one usually just considers the minimum DBI and maximum CHI over a range of possible *k*s. We have recently advocated to accompany this global analysis with a local counterpart that considers the relative stability of DBI and CHI results over a reduced range of *k*s. For example, the local stability of a partition with k clusters should be gauged against the partitions with *k* – 1 and *k* + 1 clusters. This essentially amounts to identifying local extrema in the DBI or CHI vs. *k* plots by approximating the 2^nd^ derivatives of these curves using a simple finite-differences approximation. In simple terms, we also look for the maximum value of the 2^nd^ derivative of the DBI, and the minimum value of the 2^nd^ derivative of the CHI. The local analysis helps overcoming the reported bias of these indices to prefer very low *k* values in the global case.

**Figure 4:**
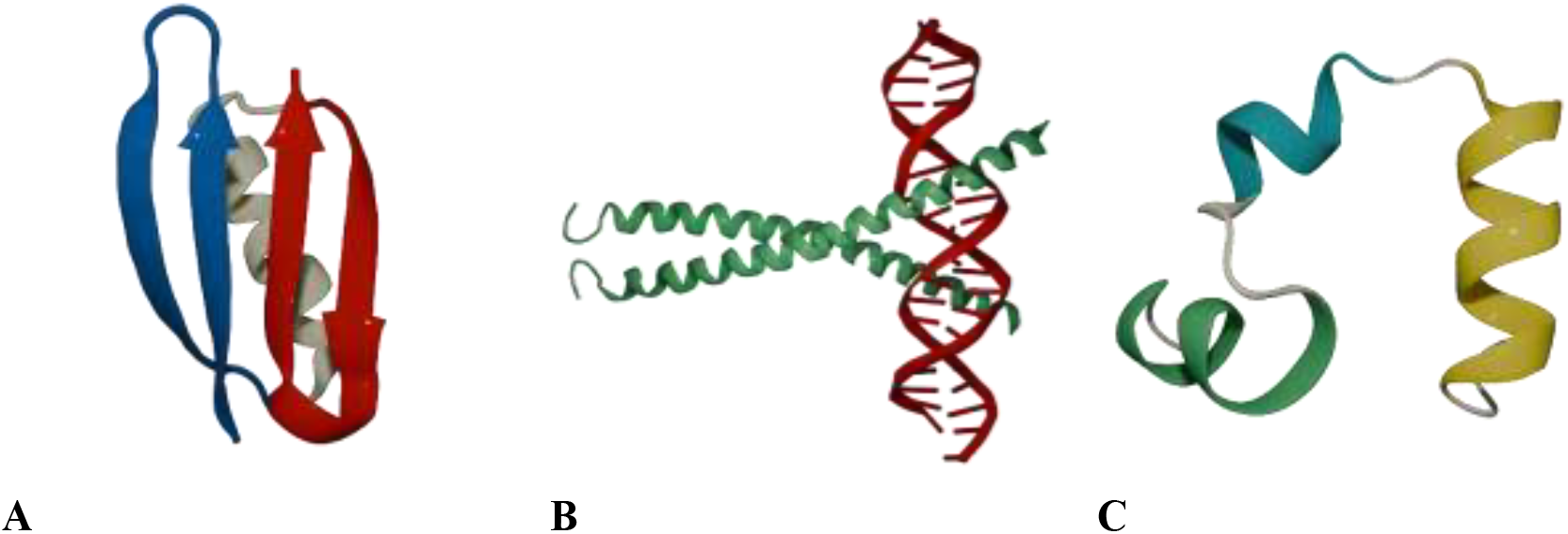
Studied systems (A) protein G, (B) bZIP binding, (C) HP35, villin headpiece. The color scheme represents the different functional regions within each system: in A, the N-termini hairpin (blue), C-termini hairpin (red), and helix (cream); in B, the helical dimer (green) and DNA (red); and in C, the three helix fragments (helix 1: green, helix 2: blue, helix 3: yellow).

## 4. RESULTS & DISCUSSION

Our first test concerns the impact on choosing different *k* values at the time of performing the NANI clustering. Fig. 5 shows the behavior of the DBI for all the studied systems when we start from 30 and 100 NANI clusters. In these cases, the 4th HELM step only involved building the hierarchy using the intra or inter linkage criteria. It is reassuring to see that, despite the drastically different initial conditions, the overall behavior in the 2-30 region is quite consistent for every system. Although minor variations in the DBI values are observed, especially in the small cluster number regions, they generally lead to local minimum within similar ranges of *k*. This suggests that the final hierarchical steps in HELM are unaffected by how finely or coarsely we initially group the data with NANI. Based on these observations, we opted to use 60 initial clusters for further analysis.

**Figure 5:**
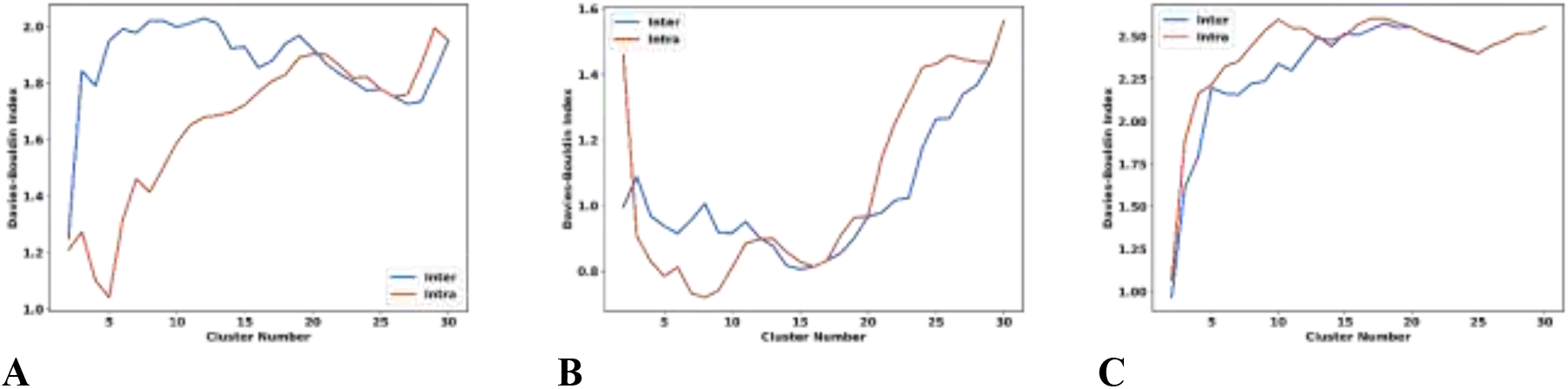
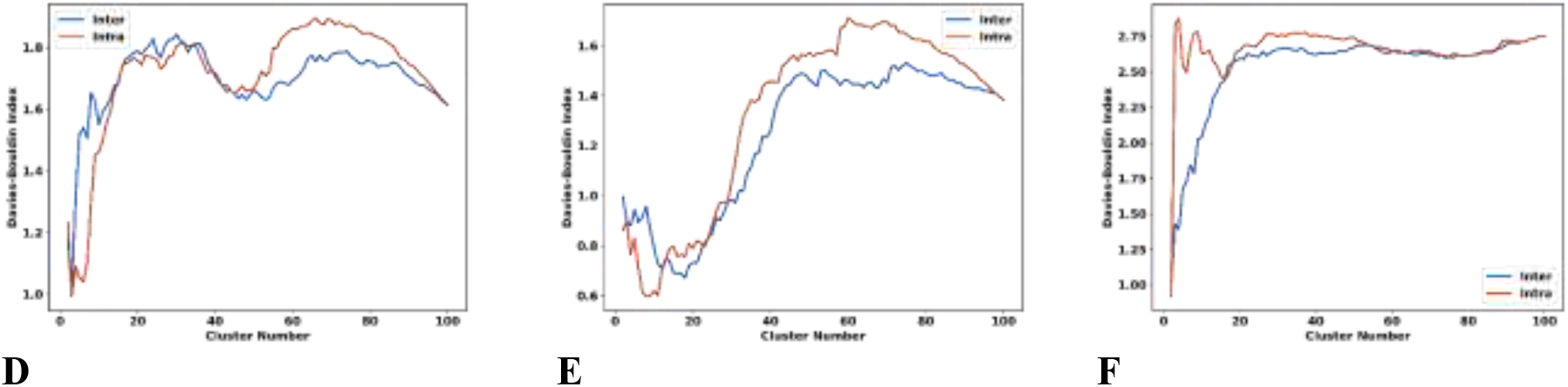
Davies-Bouldin index change for: A, D: protein G, B, E: protein-DNA, and C, F: HP35 during the hierarchical steps of HELM (*intra*: red, *inter*: blue) after starting from 30 (A, B, C) and 100 (D, E, F) NANI clusters. Another important test is how the different merging criteria in Step 4 compared to each other (starting from 60 NANI clusters, see Fig. 6). For this, we compare the intra and inter criteria with the more traditional Ward linkage. Note that we apply Ward to *k*x*k* matrices built both from the intra and inter metrics. Remarkably, the four linkages are very consistent, especially in the early stages of building the hierarchy (for numbers of clusters between 60 and 30 or 20). Also, we see the already reported tendency of the DBI and CHI to be biased towards very low cluster counts, with rapid drops (DBI) or increases (CHI) when there are only 2 or 3 clusters.

The DBI and CHI values are not only relevant to compare different clustering methods, but as noted elsewhere, they can also be used to identify particularly stable numbers of clusters in the data. As noted above, there are two different ways of doing this: a) Global analysis: just looking at either the global minimum (DBI) or maximum (CHI) over all the considered cluster counts; b) Local analysis: calculating the 2^nd^ derivative of these indices using finite differences as a way to determine which *k* values are markedly more stable than their neighbors. Also, to avoid the known bias of these indices towards low cluster counts, we restrict this type of analysis to instances with at least 5 clusters. Table 1 contains a detailed account for all the studied systems, starting from 60 NANI clusters, and using the intra, inter, Ward-intra, and Ward-inter linkages to build the hierarchy. The analysis of global DBI minima or CHI maxima, as well as their 2^nd^ derivatives, reveals how sensitive these measures can be across different linkage criteria. Given the previously reported behavior for the protein G, protein-DNA and HP35 simulations, it seems like the DBI (both absolute and 2^nd^ derivative) values do a slightly better job at identifying the proper number of clusters in the data, which is consistent with previous reports about this index.

**Table 1:** Preferred numbers of clusters for protein G, protein-DNA, and HP35.

| Linkage/System | protein G | protein-DNA | HP35 |
| --- | --- | --- | --- |
| **Minimum DBI values** |  |  |  |
| *inter* | 7 | 14 | 5 |
| *intra* | 5 | 9 | 5 |
| *inter-Ward* | 39 | 19 | 6 |
| *intra-Ward* | 37 | 15 | 5 |
| **Maximum 2<sup>nd</sup> Derivative of DBI values** |  |  |  |
| *inter* | 7 | 8 | 13 |
| *intra* | 37 | 21 | 7 |
| *inter-Ward* | 10 | 10 | 6 |
| *intra-Ward* | 7 | 31 | 7 |
| **Maximum CHI values** |  |  |  |
| *inter* | 36 | 17 | 5 |
| *intra* | 5 | 7 | 8 |
| inter-Ward | 36 | 12 | 5 |
| intra-Ward | 31 | 7 | 5 |
| **Minimum 2<sup>nd</sup> Derivative of CHI values** |  |  |  |
| inter | 26 | 9 | 36 |
| intra | 28 | 17 | 27 |
| inter-Ward | 9 | 9 | 49 |
| intra-Ward | 31 | 16 | 49 |

The results shown in Table 1 show some variability in the preferred number of clusters, but they also hint to a key issue: the impact that noisy clusters have in the overall process. For example, in Table 2 we explore the populations and MSD values for the HP35 simulation when the hierarchy gets to 6 clusters. Note that in most cases the previous clusters have just merged into an ultra-large super-cluster with over 80% of the total frames. Interestingly, the smaller clusters often exhibit substantially higher MSD values compared to the “super-cluster”, hinting that these grouped conformations are less compact. This pattern highlights that, with the inherent noise in starting data, hierarchical agglomerative clustering tends to lump most of the conformations into a broad, low-MSD cluster and only isolates some into smaller and less compact groups. This behavior can also be observed in other systems. For instance, Fig. 7 shows the overlaps of some of the conformations found in protein G and HP35 when there are 6 clusters.

**Table 2:**
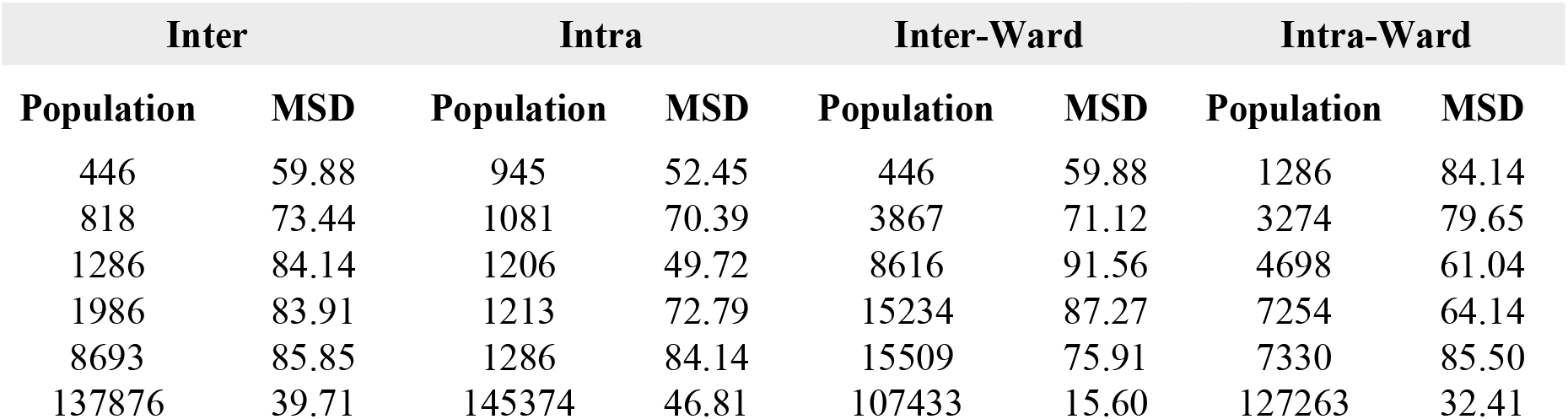
Cluster populations and MSD for *inter, intra*, inter-Ward, and intra-Ward for HP35 with *k* = 6 clusters. Clusters are ordered in increasing population.

| **Inter** |  | **Intra** |  | **Inter-Ward** |  | **Intra-Ward** |  |
| --- | --- | --- | --- | --- | --- | --- | --- |
| **Population** | **MSD** | **Population** | **MSD** | **Population** | **MSD** | **Population** | **MSD** |
| 446 | 59.88 | 945 | 52.45 | 446 | 59.88 | 1286 | 84.14 |
| 818 | 73.44 | 1081 | 70.39 | 3867 | 71.12 | 3274 | 79.65 |
| 1286 | 84.14 | 1206 | 49.72 | 8616 | 91.56 | 4698 | 61.04 |
| 1986 | 83.91 | 1213 | 72.79 | 15234 | 87.27 | 7254 | 64.14 |
| 8693 | 85.85 | 1286 | 84.14 | 15509 | 75.91 | 7330 | 85.50 |
| 137876 | 39.71 | 145374 | 46.81 | 107433 | 15.60 | 127263 | 32.41 |

**Figure 6:**
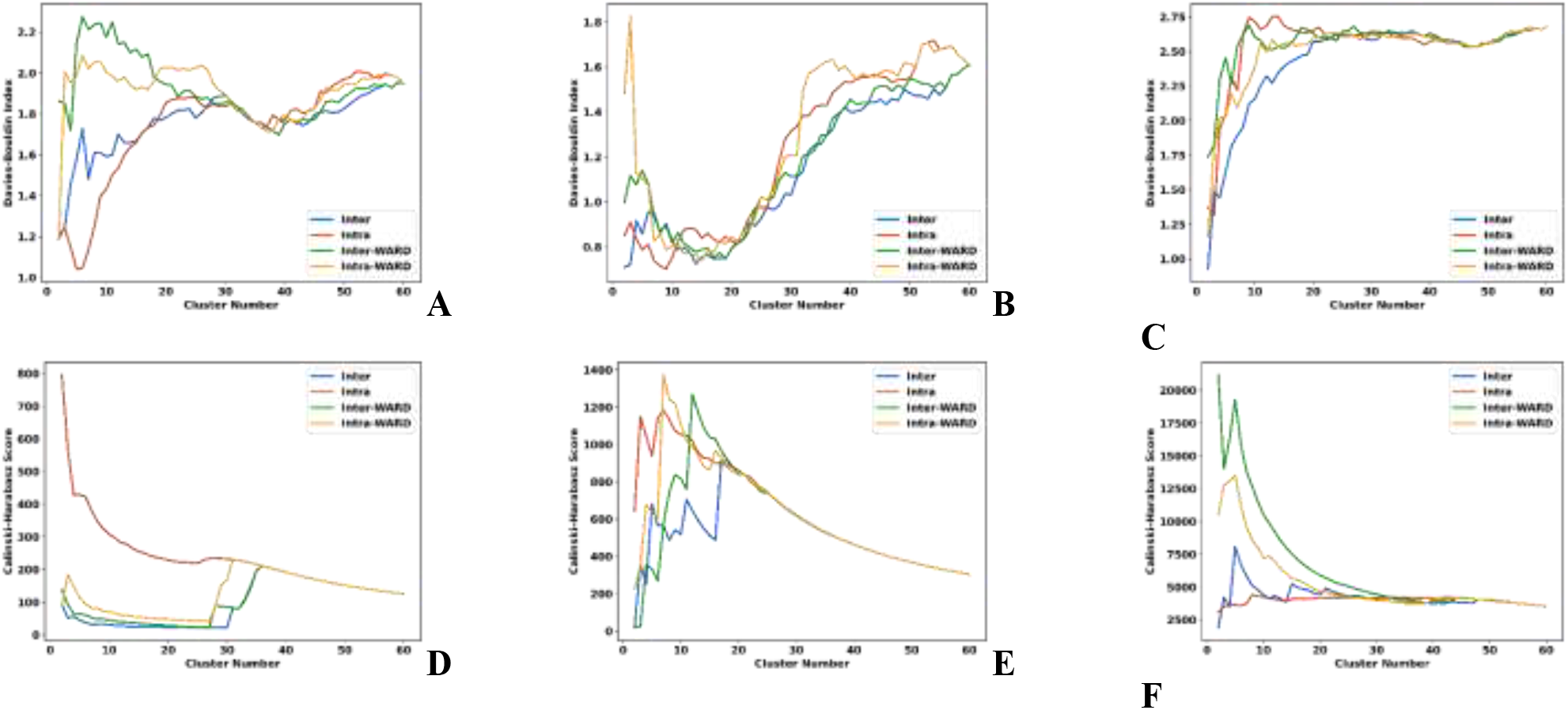
Change in Davies-Bouldin (A, B, C) and Calinski-Harabasz (D, E, F) indices for A, D: protein G, B, E: protein-DNA, and C, F: HP35 during the hierarchical steps of HELM (*intra*: red, *inter*: blue, inter-Ward: green, intra-Ward: yellow) after starting from 60 NANI clusters.

**Figure 7:**
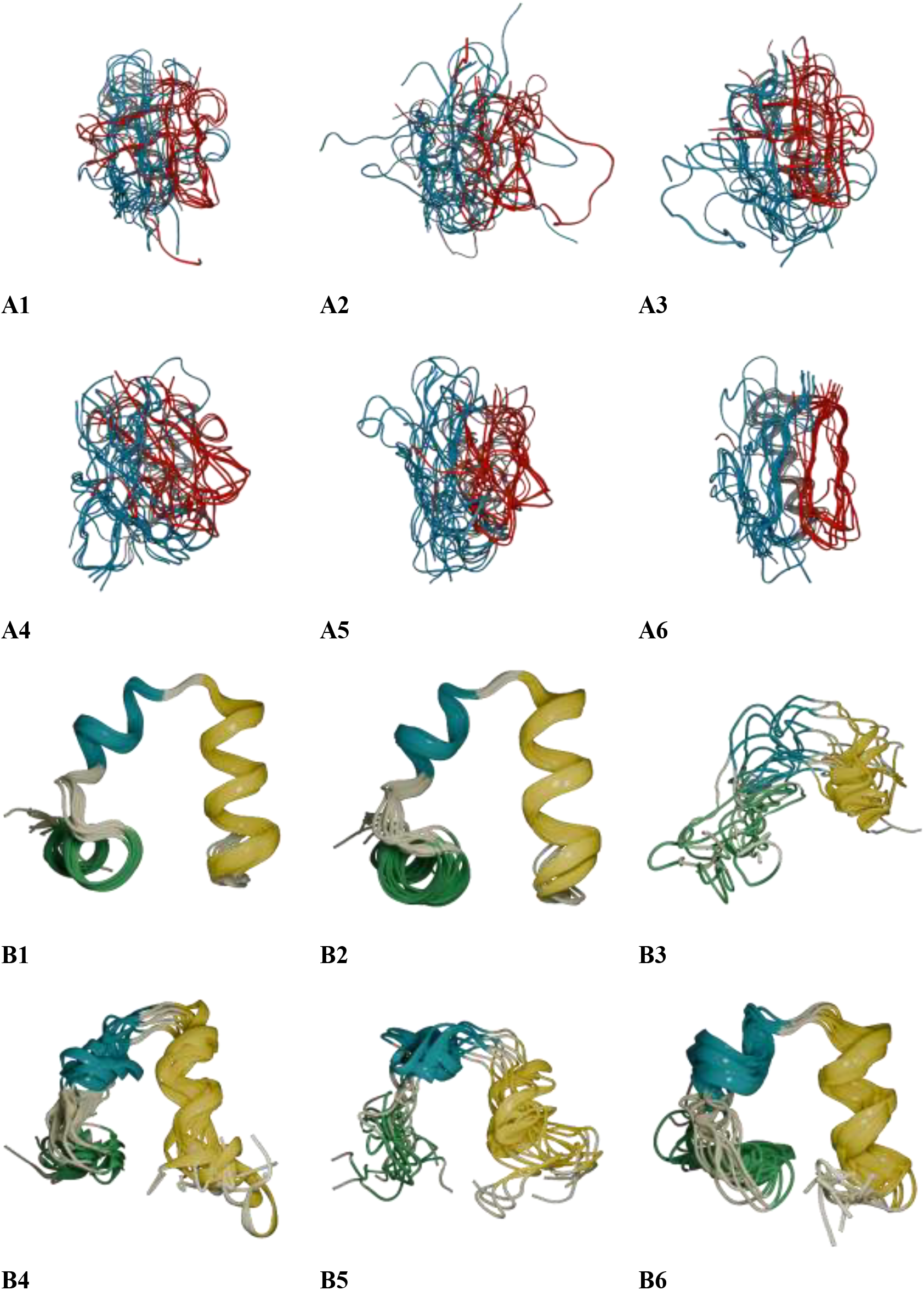
Overlaps of six clusters of the protein G (A panels) and HP35 (B panels) after performing HELM from 60 NANI clusters using the *intra* merge.

To explore how noisy clusters impact the overall hierarchical process, we consider trimming the initial set of clusters based on their MSD values and relative populations. We attempted to concentrate on the regions of the conformational landscape that had been more frequently sampled, since for some systems we observed that some of the 60 NANI clusters were essentially singletons, which means that the *k*-means pre-clustering was identifying those sectors as noisy. For instance, in protein G, the trimming process that retains clusters with MSD < 10 and at least 0.5% of the population leaves 83.57% of the frames and yields 15 clusters ready for HELM. In the protein-DNA system, the same cutoff leaves 70.25% of the data with 18 clusters remaining. In the HP35 system, we investigated two different cutoffs: Strategy A, MSD < 10, the data is trimmed to 58.70% of the population, with 14 initial clusters; Strategy B, MSD < 20, the percentage of frames increased to 66.99%, with 19 clusters. Fig. 8 shows the DBI and CHI behavior of the HP35 system when we apply trimming. Despite the different thresholds, most of the trends are consistent and the DBI minima and CHI maxima fall within the similarly low-number cluster regions. Interestingly, for MSD < 20 trimming, the inter linkage exhibits a sharp decrease of CHI when going from six to five clusters, a trend not seen in the plot of the intra linkage. Abrupt changes like this may be used as indicators that merging those clusters may lower the overall quality of the clustering, which indicates an optimal *k* value.

**Figure 8:**
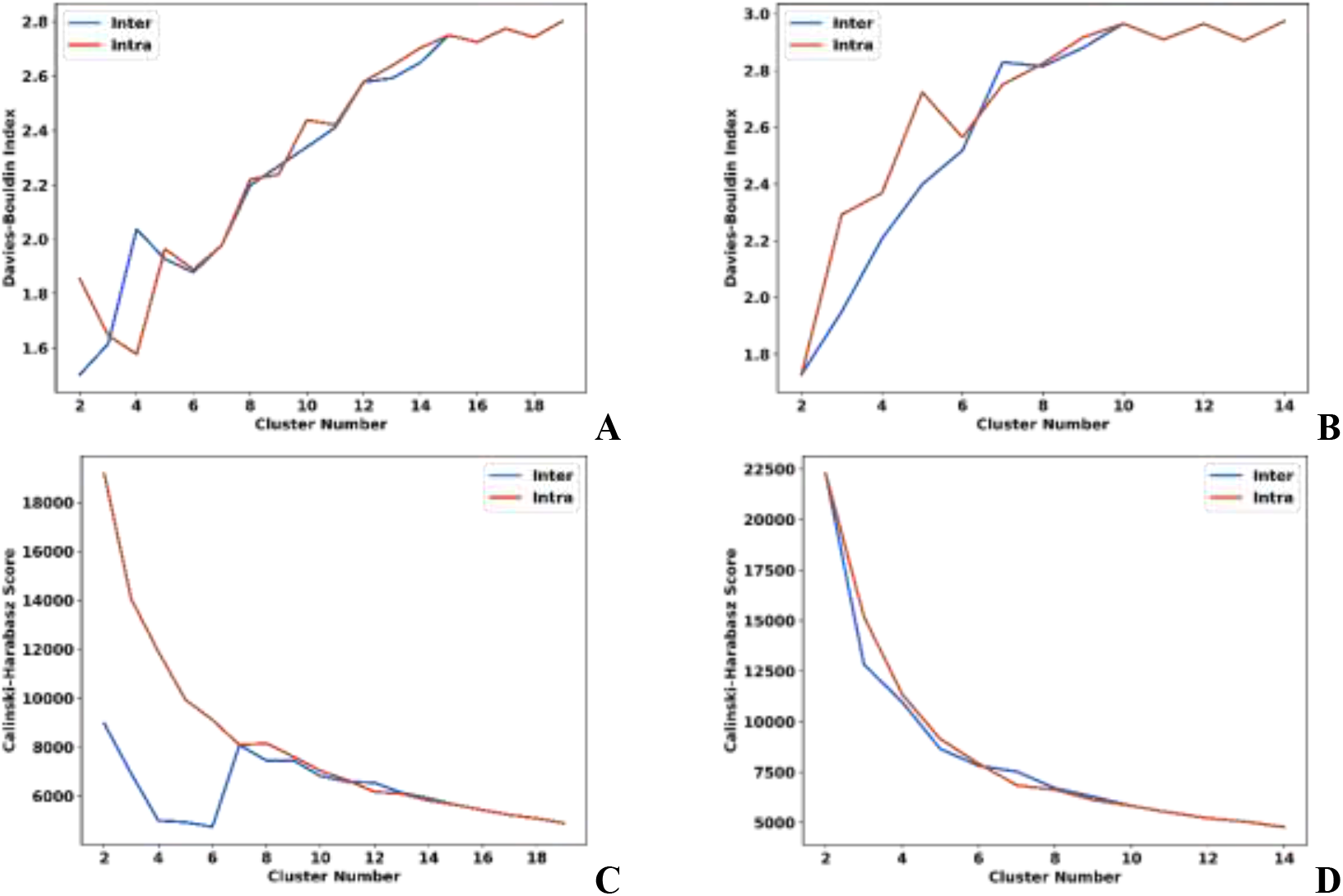
Variation in the Davies-Bouldin (A, B) and Calinski-Harabasz (C, D) indices for the HP35 simulation after trimming the initial NANI clusters with MSD < 20 (A, C) and MSD < 10 (B, D). *intra* merge in red, *inter* merge in blue.

Table 3 contains the preferred number of clusters for all the studied systems, after trimming the noisy sectors, and using only the intra and inter linkages to build the hierarchy. Here we use the same global and local analysis in predicting an optimal number of clusters after the hierarchical process, while also restricting the analysis to five clusters to eliminate the mentioned low cluster counts bias. It is reassuring to see the robust behavior of all these methods given the elimination of the noisy sectors, while producing numbers of clusters in close agreement with previous studies on these systems.

**Table 3:** Preferred numbers of clusters for protein G, protein-DNA, and HP35 after trimming the noisy clusters (blank cells indicate that no local minimum met the criteria).

| Linkage/System | protein G | protein-DNA | HP35 * |
| --- | --- | --- | --- |
| **Minimum DBI values** |  |  |  |
| inter | 6 | 5 | 5/6 |
| intra | 6 | 5 | 6/6 |
| **Maximum 2<sup>nd</sup> Derivative of DBI values** |  |  |  |
| inter | 6 | 8 | 13/6 |
| intra | 6 | 11 | 6/11 |
| **Maximum CHI values** |  |  |  |
| inter | 6 | 5 | 5/7 |
| intra | 6 | 5 | 5/5 |
| **Minimum 2<sup>nd</sup> Derivative of CHI values** |  |  |  |
| inter | 6 | - | -/9 |
| intra | 6 | - | -/8 |

\* MSD < 10 / MSD < 20

Comparing the data in Table 3 from the earlier results in Table 1, notable shifts are present in the preferred cluster counts when we apply the trimming criteria. After the removal of high-MSD and small-sized clusters for the HELM process, the optimal cluster counts converge to relatively small and more consistent values across all linkages and systems. This suggests that the “reduction” of noise by trimming leads to a more compact and clean partitioning of the dataset. These observations arise because trimming removes the high-MSD clusters that would otherwise be merged with other relatively more stable clusters, inflating their MSD as they contribute to small number of frames. Ultimately, the exclusion of the noise greatly improves the resulting hierarchy, since now the merging process is not influenced by the presence of outliers that were helping make artificial connections between the clusters.

Fig. 9 shows the overlaps of the conformations found in the protein G and HP35 when there are 6 clusters (after intra linkage clustering and trimming: MSD < 10). For protein G, the secondary structure elements generally remain aligned across the clusters, but there are differences in the orientations of the loops making each cluster distinct. In HP35, the clusters also exhibit similar α-helix regions, but they vary on how the helices turn and the tails arrange themselves. Table 4 presents the cluster populations and MSD values after applying trimming. In contrast to the previous run without trimming, where only the largest population exhibited a low MSD, the trimmed results show consistently lower MSD values. Eliminating “noisy” clusters that contain mostly outlier conformations allowed HELM to focus on the core structural variations present in the system. The improved distribution of the population also helps in ensuring that no single cluster dominates, as was the case before trimming. Moreover, these populations also reflect the behavior reported for this system from a purely NANI study and a more elaborate shape-GMM analysis.

**Table 4:**
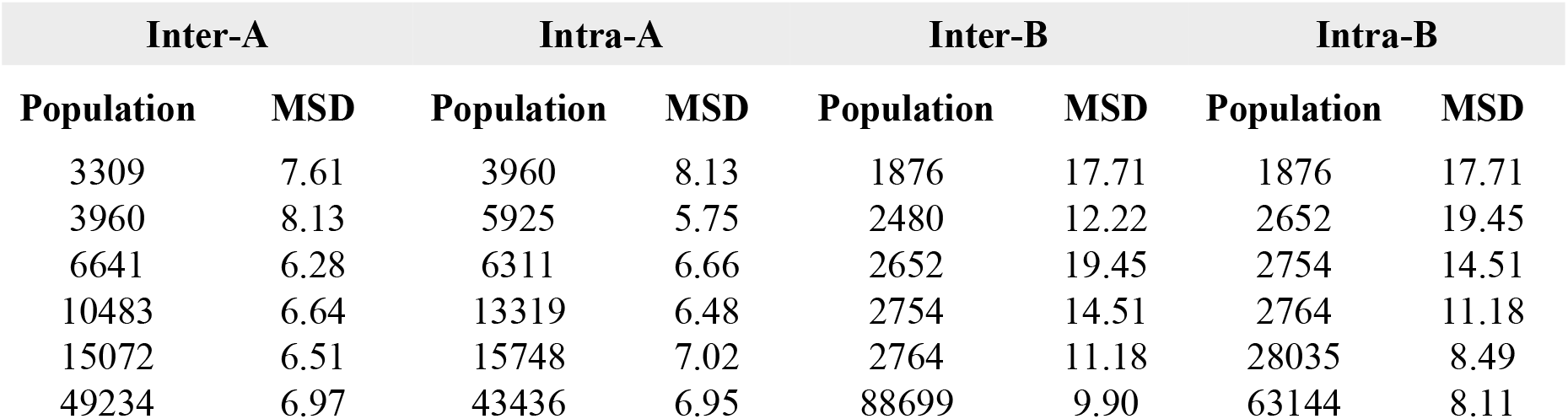
Cluster populations and MSD for *inter* and *intra* merging for HP35 *k* = 6 after trimming the noisy clusters. Clusters are ordered in increasing population.

**Figure 9:**
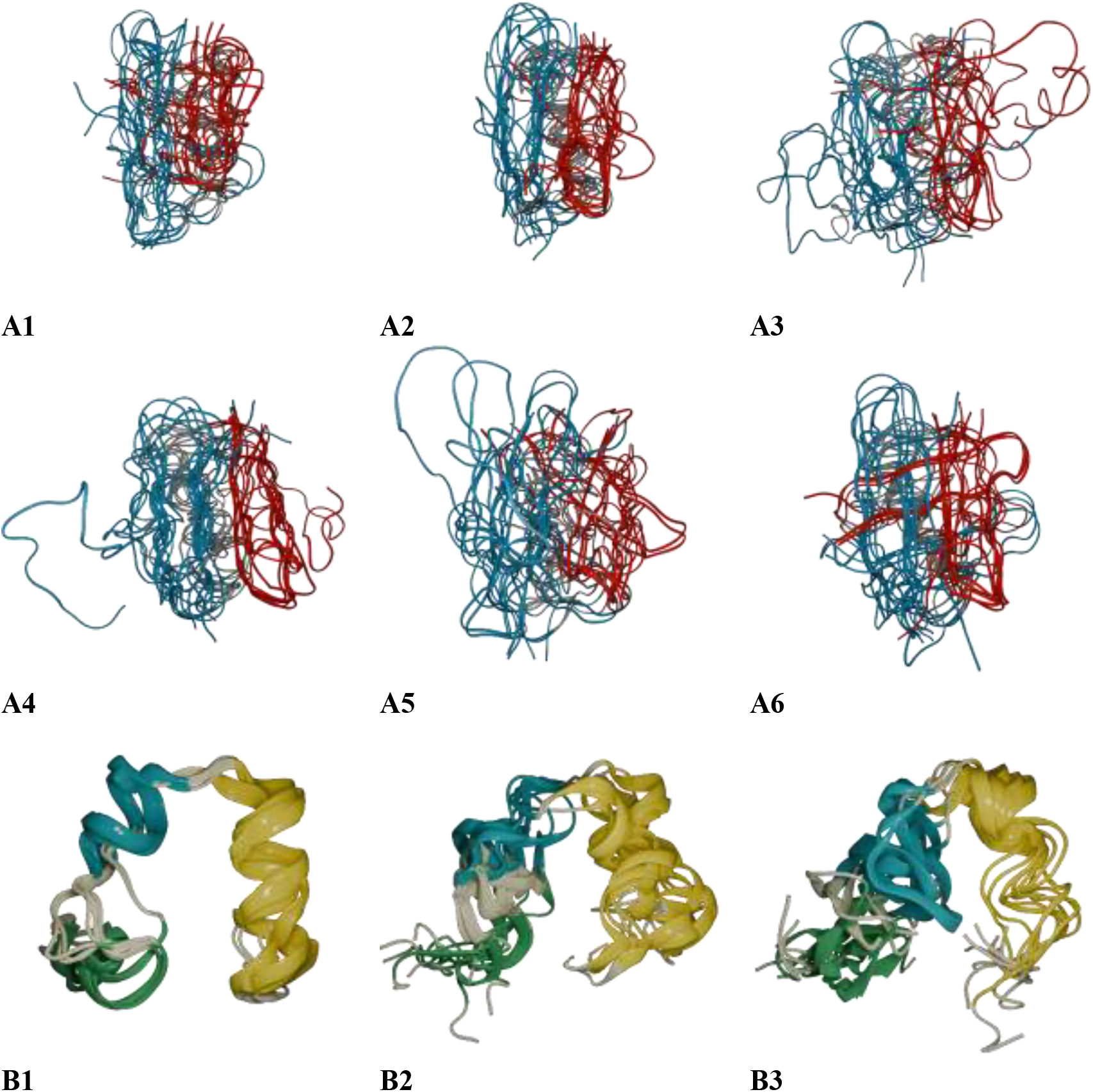
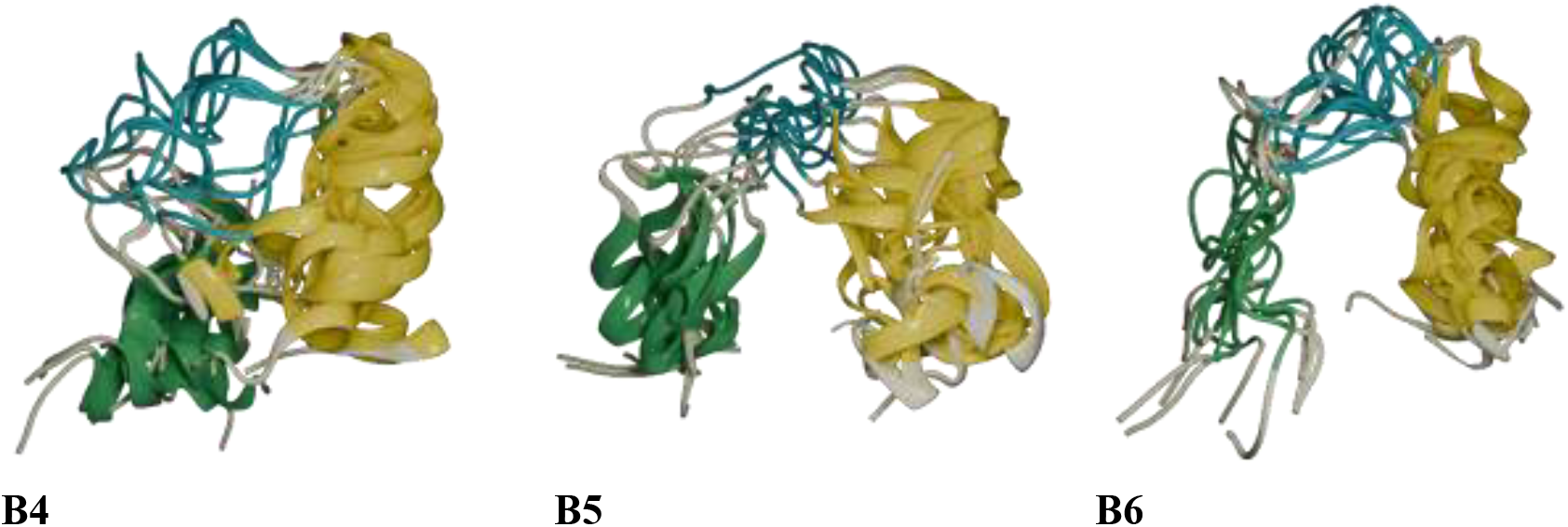
Overlaps of six clusters of the protein G (A panels) and HP35 (B panels) after performing HELM from 60 NANI clusters using the *intra* merge after the noisy clusters are trimmed.

Up to this point, we have mostly been concerned with the quality of the HELM clusters, but now we shift our attention to one of the critical issues mentioned at the beginning of this manuscript: the computational efficiency of traditional hierarchical approaches. As noted above, existing HAC methods scale quadratically in time, making it a great burden when clustering very long simulations. Table 5 demonstrates the significant improvements of HELM-based clustering over conventional methods, specifically Average and Single linkages using CPPTRAJ^44^ using Amber 22^45^(which is remarkable, given the extremely efficient implementations of these methods in this package). For the protein G and protein-DNA systems, our method completes the clustering of the systems almost twice as fast as the conventional approaches. This alone shows a notable increase in efficiency, even considering that these are very short simulations.

**Table 5:**
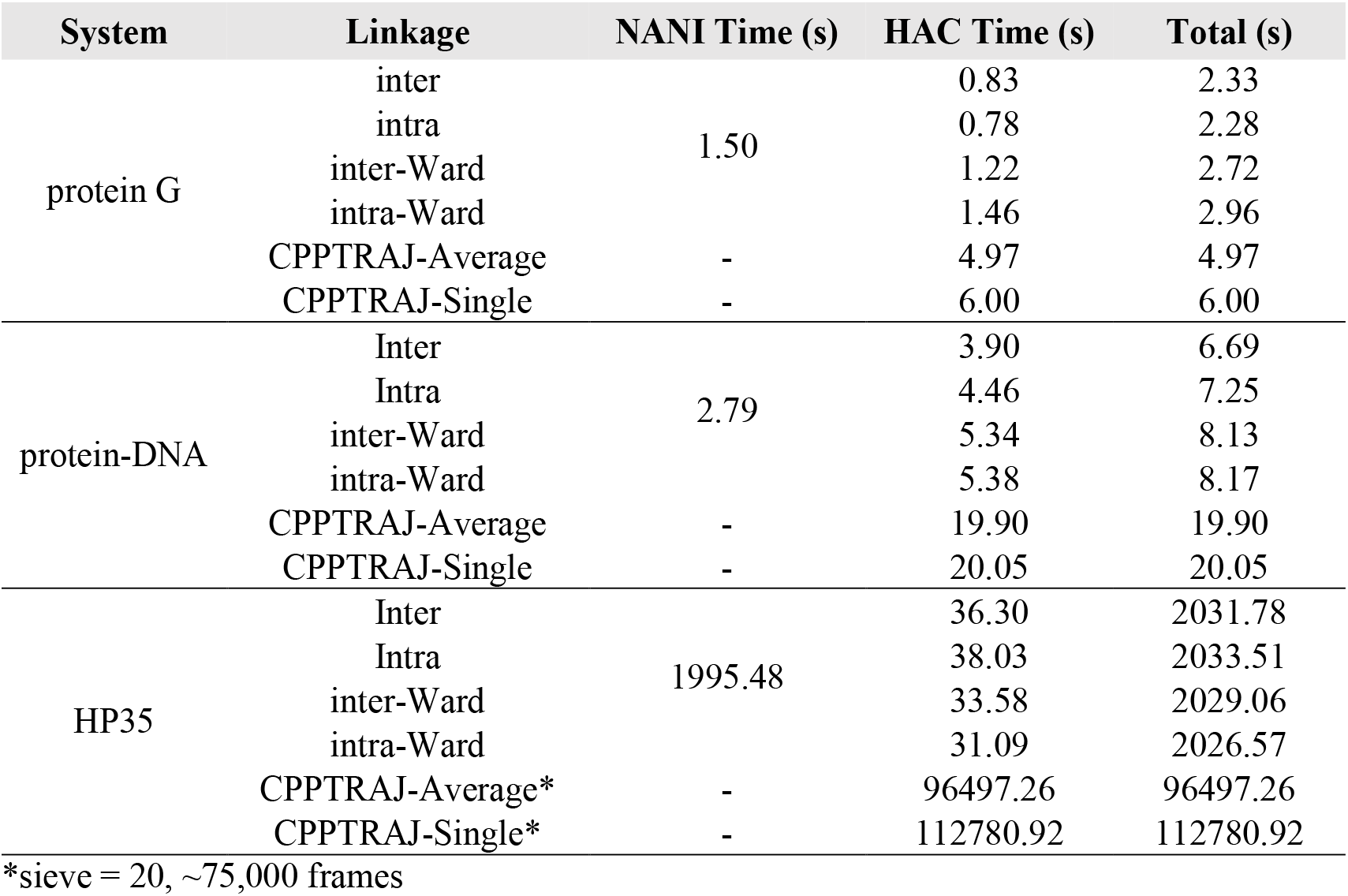
Times for the HELM and CPPTRAJ processing of protein G, protein-DNA and HP35 (with a sieve of 10). HELM starts with 60 clusters.

However, we can see the most extreme improvement in the HP35 system, where HELM completes the clustering of ∼150,000 conformations in 34 minutes, while the Average and Single methods require 26.8 and 31.3 hours, just to cluster ∼75,000 conformations, respectively. This speedup demonstrates that HELM-based clustering is much more well-positioned to handling very large datasets. As an example of this, we used HELM to cluster the full ∼ 1.5 million frames of the HP35 trajectory made public by D. E. Shaw Research (to the best of our knowledge, the first hierarchical clustering of this full trajectory), which only required 29 hours. Following the trends shown in Fig. 2, we estimate that completing the same analysis with CPPTRAJ will require 52 days (Average HAC) or 258 days (Single HAC). The key factor in HELM’s efficiency is the NANI step (Step 1), which pre-clusters the data before the hierarchical merging. While this step is much costlier than building the final hierarchy, this is the key responsible for speeding up Steps 2 to 4, when compared to the CPPTRAJ methods.

## 5. CONCLUSIONS

We have presented HELM as a hybrid strategy combining the time and memory efficiency of *k*-means with the flexibility of hierarchical approaches. The natural way of combining these methods possess the question: why was this hybrid framework not explored in more detail before? The short answer is that to be able to truly take advantage of all the *k*-means and HAC features, it was necessary to have an O(*N*) way to quantify similarity, which was not possible until the introduction of iSIM or the MSD. In more detail, from the *k*-means part, it is important to have a fully deterministic (reproducible) starting pre-clustering step, since this is the only way to have a stable hierarchy. MSD makes this possible thanks to *k*-means NANI, which is not only a robust *k*-means recipe, but also one that does not compromise on the quality of the *k*-means segmentation. Then, for the HAC part, we need to be able to easily quantify the separation between sets of conformations without needing the individual information of every single frame in each cluster. This is not possible with only the RMSD, since by construction it can only operate over well-defined pairs of individual frames. Once again, the MSD overcomes this need by providing natural ways to determine the relation between sets, as reflected in the *intra* and *inter* merging protocols. The application of HELM to multiple systems highlights its ability to capture the underlying structure of the conformational landscape of realistic MD simulations. As was the case for the pure NANI studies, the combination of the global and local DBI and CHI analyses helps identifying optimal partitions of the data, which reassuringly agree with previous studies on these systems. Moreover, we showed how the NANI step can be used to identify low density, noise, and subsets in the data, which after being excluded from the hierarchy greatly improve the final clustering results. This is traditionally not possible in a pure *k*-means study, since this method has no notion of “noise”, and every single point will be assigned to every possible cluster. Given the relatively small number of clusters usually obtained after a full *k*-means study this translates into much of the noise being distributed over these clusters, which complicates identifying subsets of unrelated conformations. However, in this case we do not require *k*-means to produce the final assignment, and the fact that we are purposely generating many more clusters than we anticipate having in the end makes it easier to identify the noisy data. This functionality resembles the popular feature of HAC methods of stopping hierarchy when a given threshold of inter-cluster separation is found, but with the added advantage of skipping hundreds (if not thousands) of singlet-merging steps. It is important to remark that the advantages brought forward by HELM arise from the algorithmic gains provided by the MSD, which results in a fundamentally different way of approaching the problem of building a hierarchy of clusters. For example, the CPPTRAJ HAC implementation is as optimized as it could possibly be, but since it is based on the RMSD it cannot escape the need to build a pairwise matrix of frame-to-frame distances. The sheer memory demands of this step make it virtually impossible to tackle simulations with hundreds of thousands (let alone millions) of conformations. It is remarkable that HELM, with a far less optimized implementation than those found in CPPTRAJ or scikit-learn, can still process much larger sets in a fraction of the time. In our case, as seen in Table 5, the NANI step is the time-determining bottleneck. Given the promise of both NANI and HELM, we are currently working on optimizing this implementation.

## ACKNOWLEDGEMENTS

RAMQ, LC, and JBWS thank support from the National Institute of General Medical Sciences of the National Institutes of Health under award number R35GM150620. AP, JG thank the National Science Foundation CAREER award, CHE-2235785.

