## Supplementary Information for "Hierarchical Extended Linkage Method (HELM)’s Deep Dive into Hybrid Clustering Strategies"

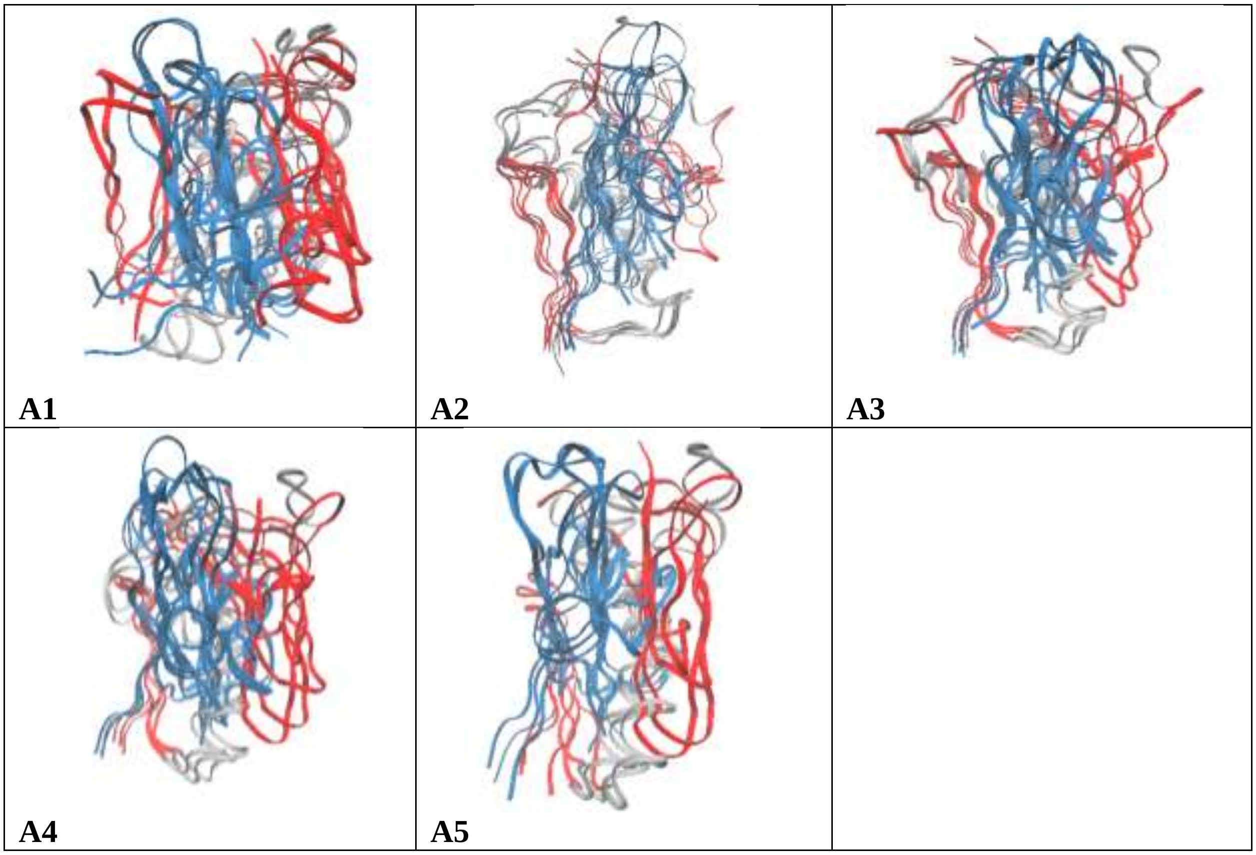

| | | |
|---|---|---|
| 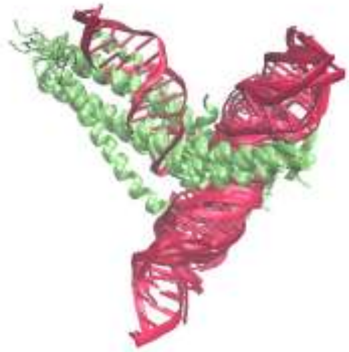 B1 | 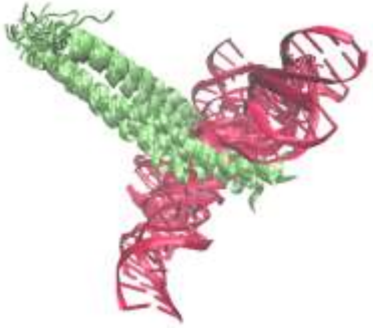 B2 | 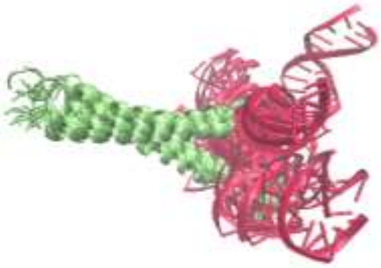 B3 |
| 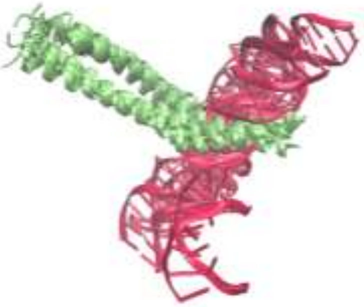 B4 | 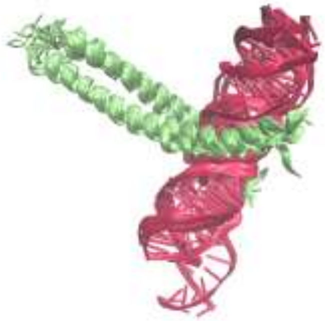 B5 | 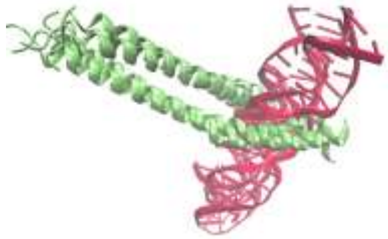 B6 |
| 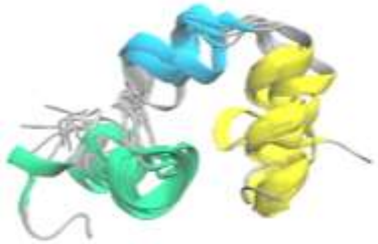 C1 | 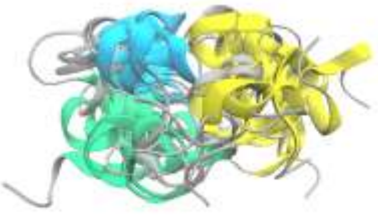 C2 | 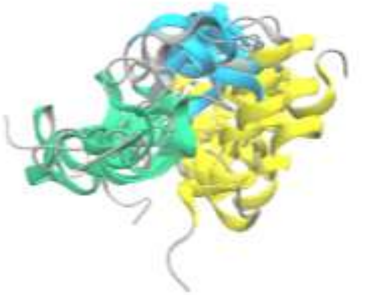 C3 |
| 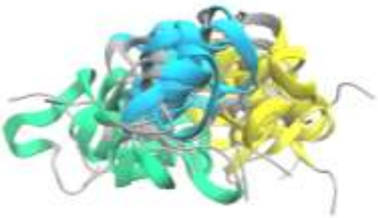 C4 | 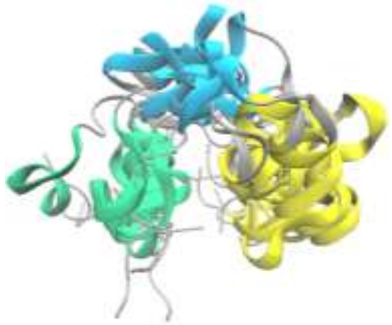 C5 | 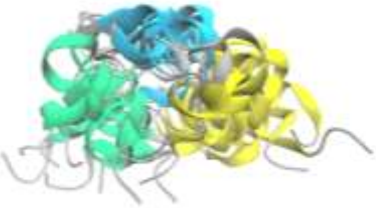 C6 |

**Figure S1:** Overlaps of six clusters of the protein G (A panels), protein-DNA binding (B panels), and HP35 (C panels) after performing HELM from 60 NANI clusters using the inter merge. Note: No image was generated for the last protein G cluster because it contained only a single frame.

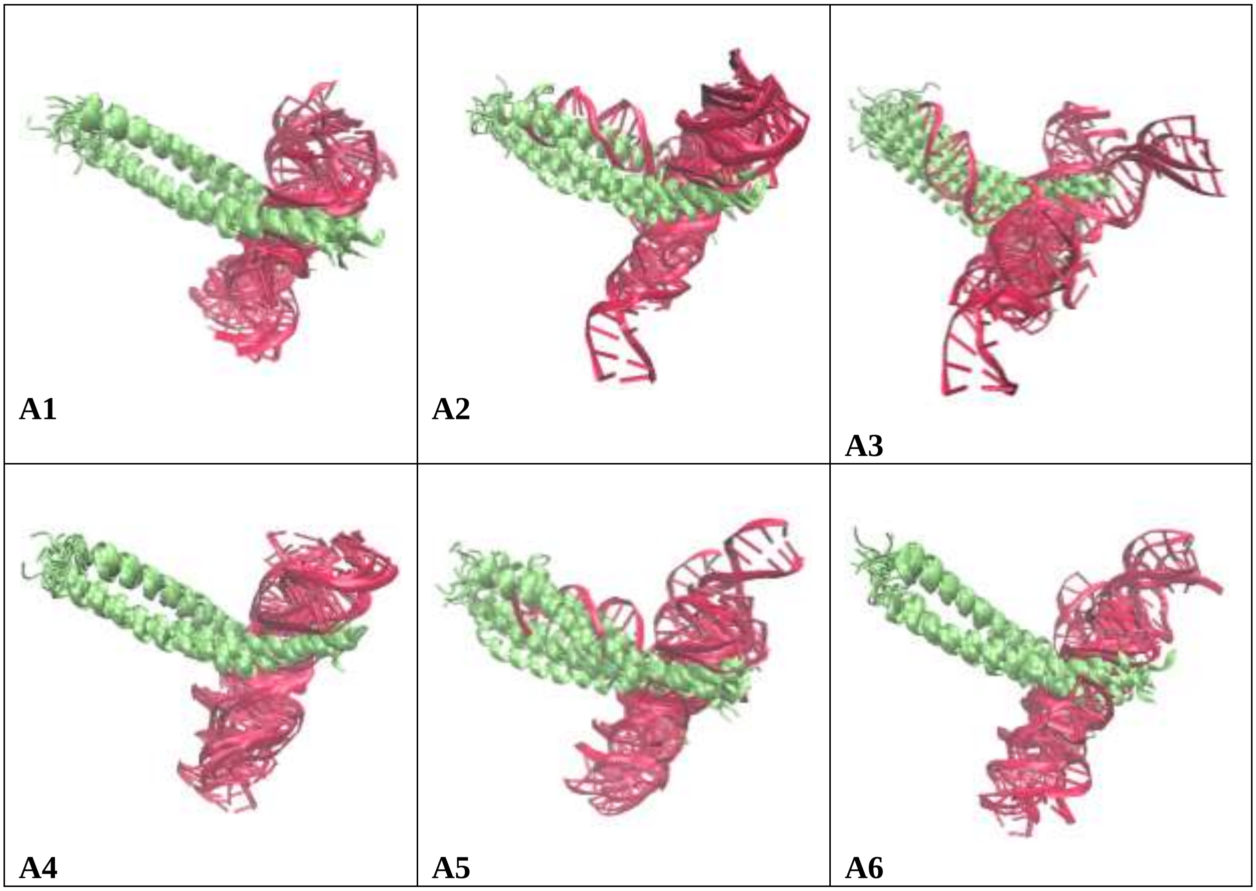


**Figure S2:** Overlaps of six clusters of the protein-DNA binding after performing HELM from 60 NANI clusters using the *intra* merge.

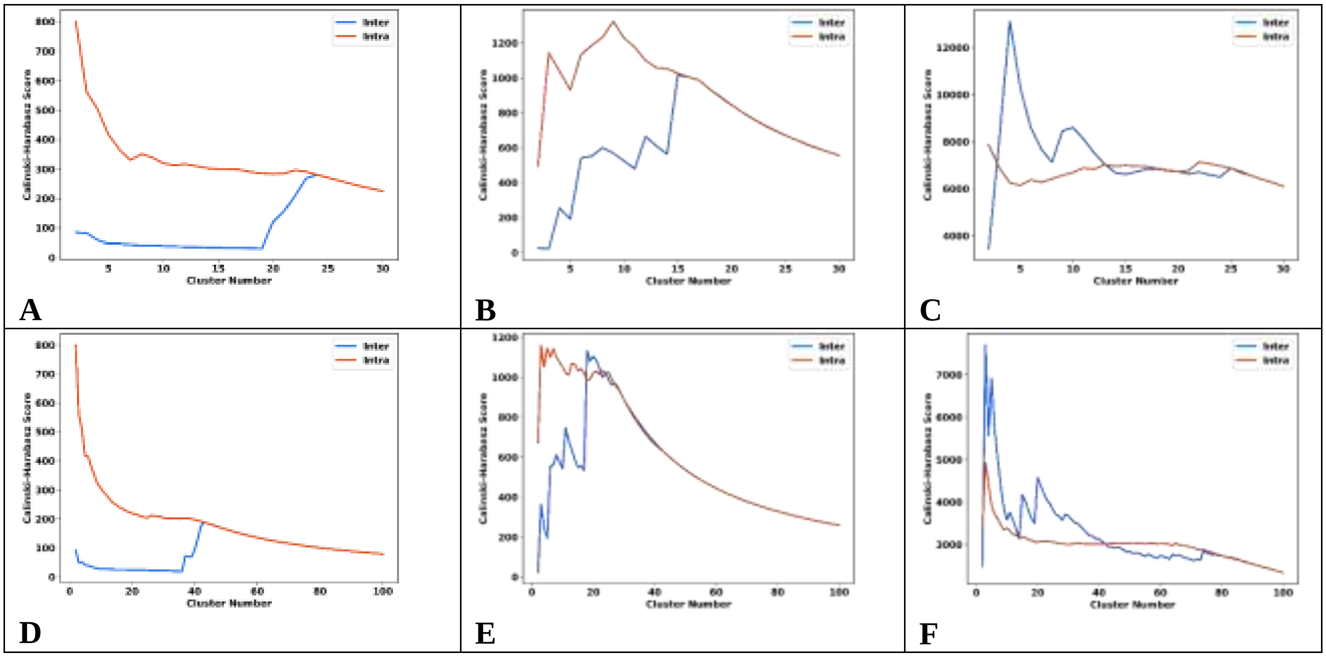


**Figure S3**: Calinski-Harabasz index change for: A, D: protein G; B, E: protein-DNA; and C, F: HP35 during the hierarchical steps of HELM (*intra*: red, *inter*: blue) after starting from 30 (A, B, C) and 100 (D, E, F) NANI clusters.

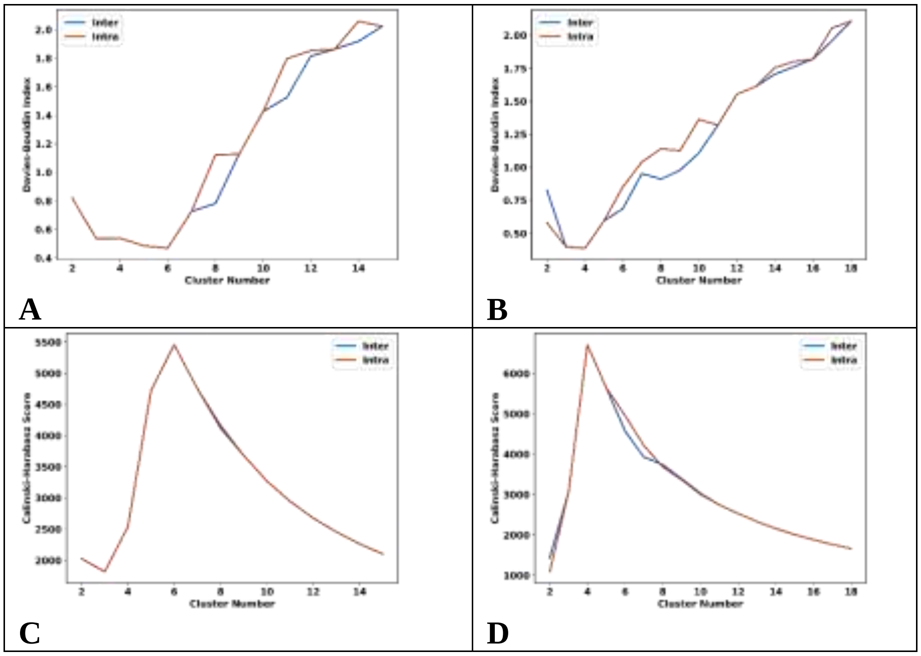


**Figure S4**: Change in Davies-Bouldin (A, B) and Calinski-Harabasz (C, D) indices for A, C: protein G and B, D: protein-DNA simulation after trimming the initial NANI clusters (retaining clusters with MSD < 10 and at least 0.5% of the population). *intra* merge in red, *inter* merge in blue.

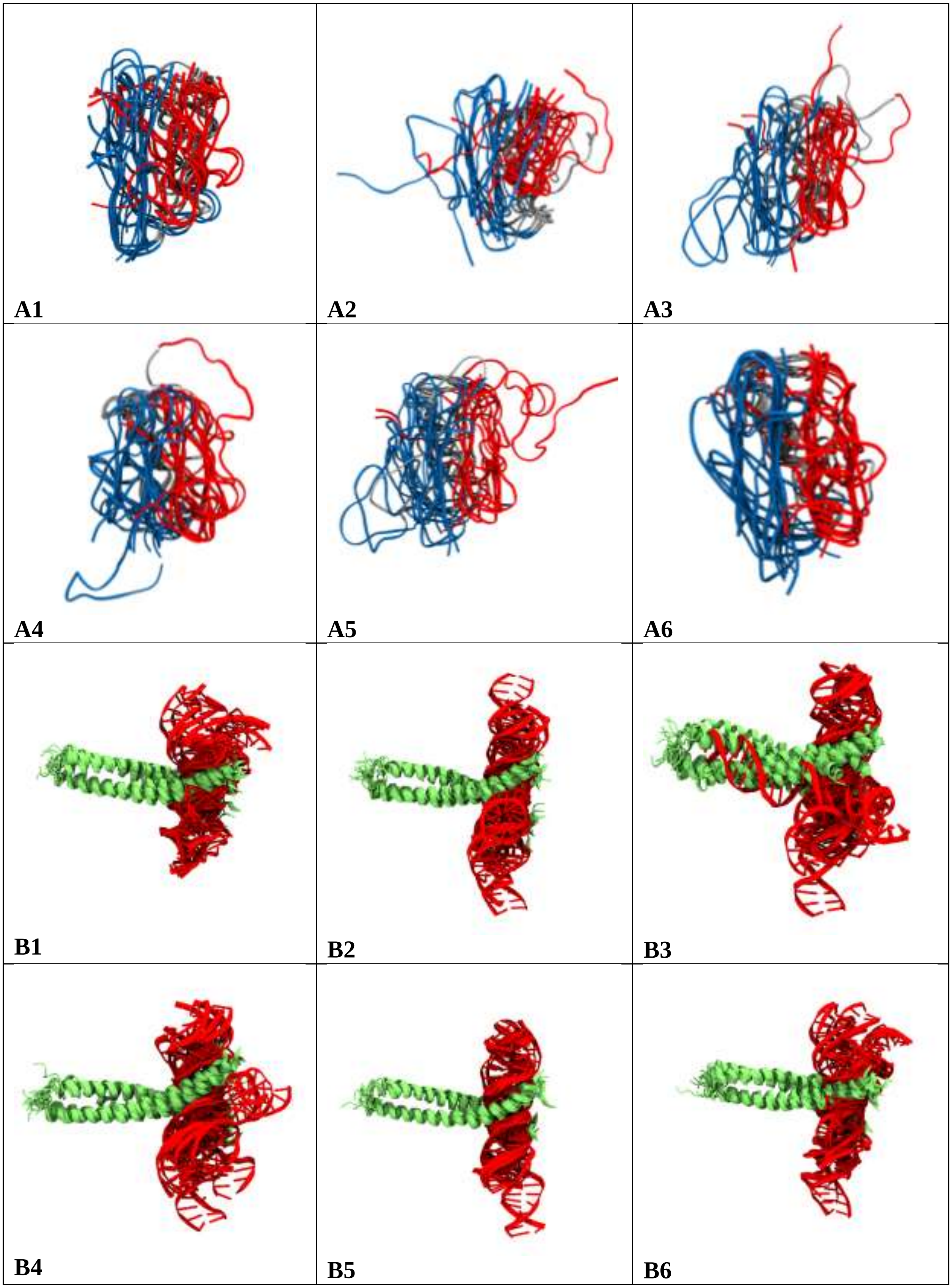
A1
A2
A3
A4
A5
A6
B1
B2
B3
B4
B5
B6

| | | |
|---|---|---|
| 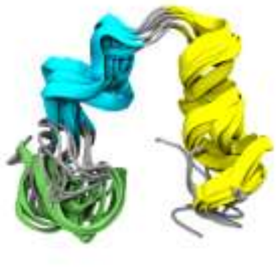 C1 | 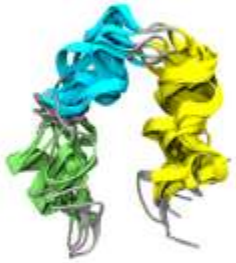 C2 | 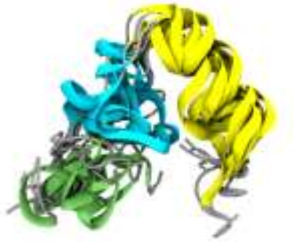 C3 |
| 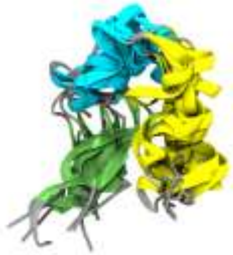 C4 | 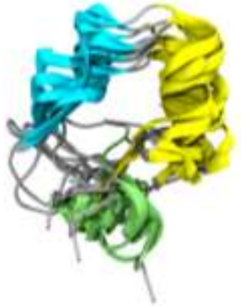 C5 | 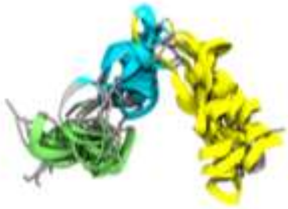 C6 |
| 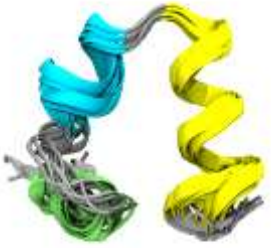 D1 | 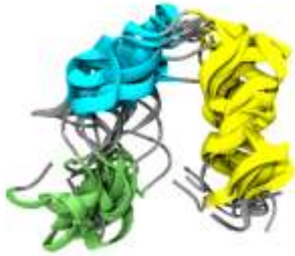 D2 | 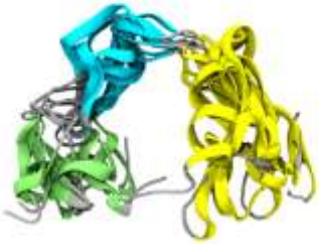 D3 |
| 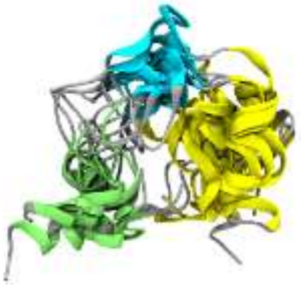 D4 | 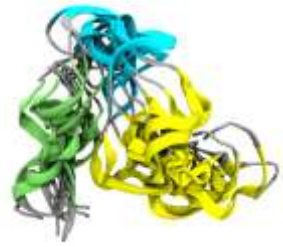 D5 | 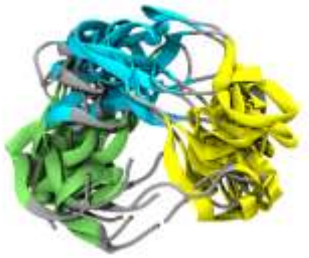 D6 |

**Figure S5:** Overlaps of six clusters of the protein G (A panels), protein-DNA binding (B panels), and HP35 (MSD < 10: C Panels, MSD < 20: D Panels) after performing HELM from 60 NANI clusters with trimming using the inter merge.

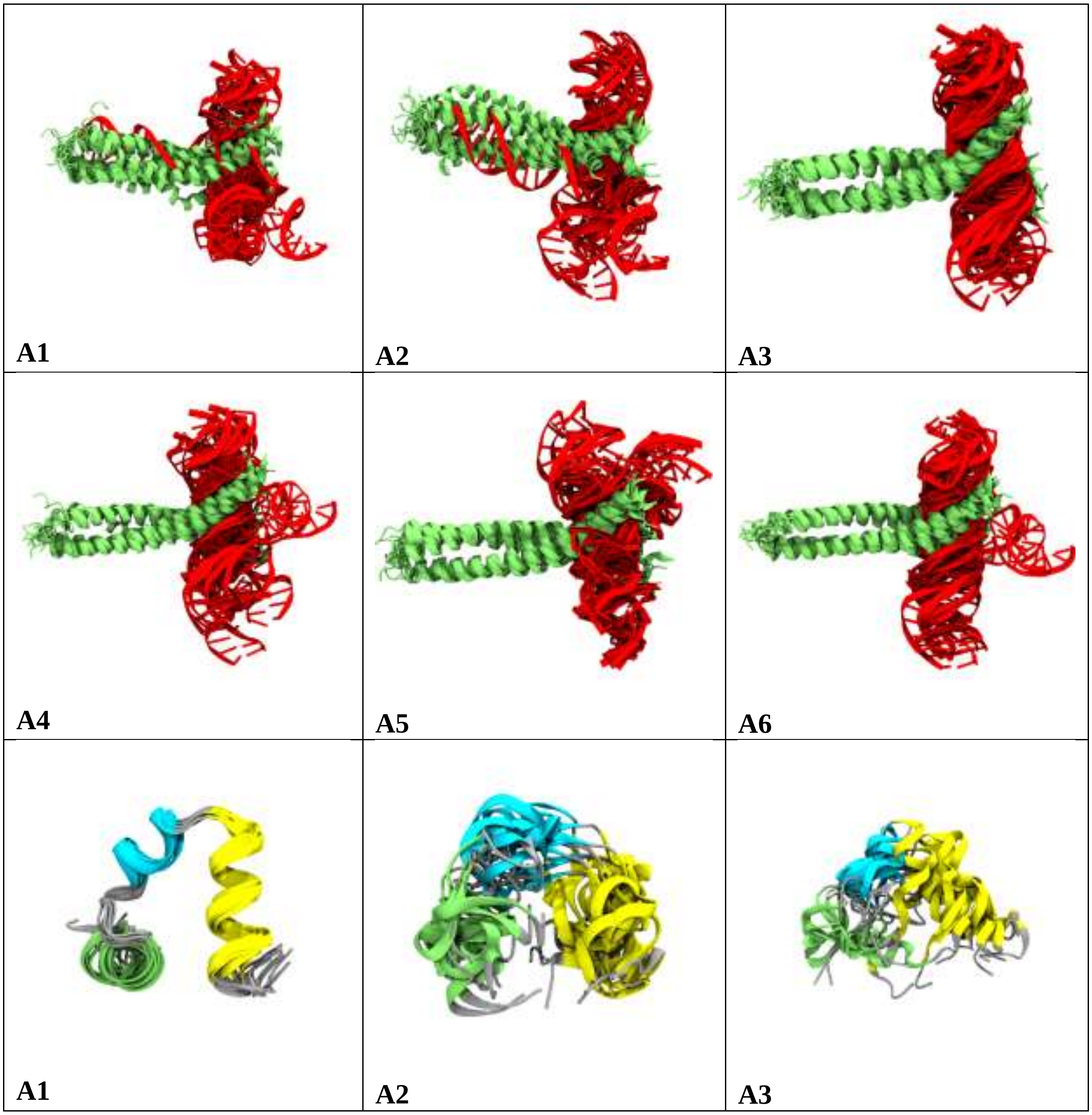

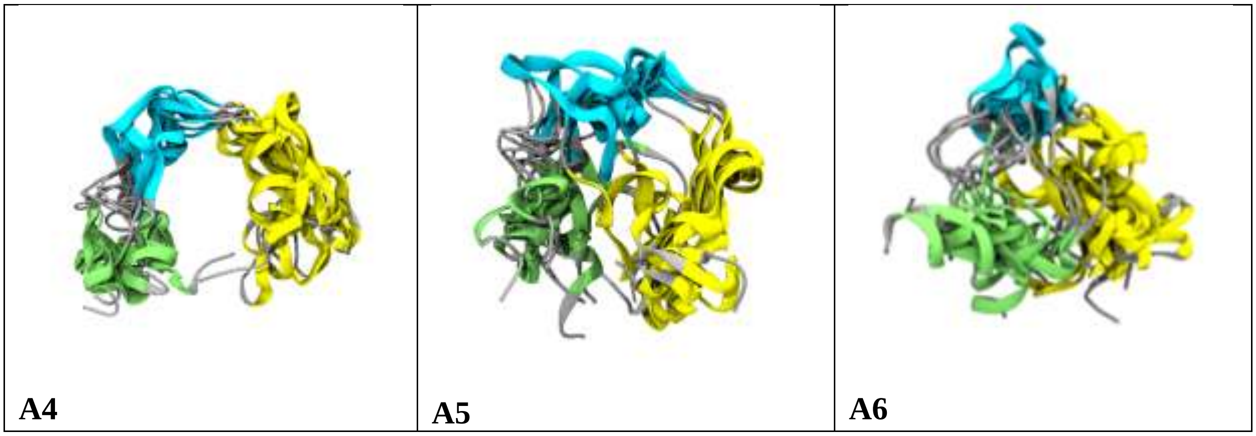


**Figure S6:** Overlaps of six clusters of the protein-DNA binding (A panels), and HP35 (MSD < 10: B Panels) after performing HELM from 60 NANI clusters with trimming using the intra merge.

**Table S1:** Preferred numbers of clusters for protein G, protein-DNA, and HP35 when starting with 30/100 NANI clusters.

| Linkage/System | protein G | protein-DNA | HP35 |
|---|---|---|---|
| | **Minimum DBI values** | | |
| inter | 27/7 | 15/18 | 7/5 |
| intra | 5/6 | 8/8 | 5/16 |
| | **Maximum 2nd Derivative of DBI values** | | |
| inter | 16/7 | 6/33 | 11/8 |
| intra | 8/6 | 29/57 | 25/6 |
| | **Maximum CHI values** | | |
| inter | 24/43 | 15/18 | 5/5 |
| intra | 5/5 | 9/5 | 22/5 |
| | **Minimum 2nd Derivative of CHI values** | | |
| inter | 15/17 | 8/27 | 18/52 |
| intra | 12/37 | 9/21 | 15/60 |

**Table S2:** Cluster populations and MSD for *inter*, *intra* for protein G with $k$ = 6 clusters with and without trimming. Clusters are ordered in increasing population.

| Inter | | Intra | | Inter with Trimming | | Intra with Trimming | |
|---|---|---|---|---|---|---|---|
| **Population** | **MSD** | **Population** | **MSD** | **Population** | **MSD** | **Population** | **MSD** |
| 2 | 111.50 | 14 | 185.31 | 176 | 1.29 | 176 | 1.29 |
| 7 | 150.78 | 250 | 38.44 | 219 | 3.98 | 219 | 3.98 |
| 9 | 174.33 | 265 | 46.49 | 237 | 9.10 | 237 | 9.10 |
| 12 | 157.57 | 281 | 49.61 | 239 | 2.50 | 239 | 2.50 |
| 16 | 188.83 | 374 | 43.57 | 340 | 3.86 | 340 | 3.86 |
| 1877 | 90.56 | 739 | 51.49 | 396 | 2.49 | 396 | 2.49 |

**Table S3:** Cluster populations and MSD for *inter*, *intra* for protein-DNA with $k$ = 6 clusters with and without trimming. Clusters are ordered in increasing population.

| Inter | | Intra | | Inter with Trimming | | Intra with Trimming | |
|---|---|---|---|---|---|---|---|
| **Population** | **MSD** | **Population** | **MSD** | **Population** | **MSD** | **Population** | **MSD** |
| 3 | 70.05 | 60 | 51.93 | 8 | 2.38 | 77 | 2.80 |
| 9 | 52.68 | 179 | 13.32 | 77 | 2.80 | 98 | 2.54 |
| 13 | 41.33 | 204 | 25.73 | 145 | 3.69 | 153 | 4.08 |
| 287 | 5.13 | 306 | 26.59 | 175 | 9.34 | 175 | 9.34 |
| 439 | 71.00 | 378 | 37.07 | 287 | 5.13 | 275 | 3.34 |
| 765 | 44.36 | 389 | 20.20 | 373 | 4.44 | 287 | 5.13 |
